# Ultrastructural analysis of nasopharyngeal epithelial cells from patients with SARS-CoV-2 infection

**DOI:** 10.1101/2021.07.08.451607

**Authors:** Karina Lidianne Alcântara Saraiva, Luydson Richardson Silva Vasconcelos, Matheus Filgueira Bezerra, Rodrigo Moraes Loyo Arcoverde, Sinval Pinto Brandão-Filho, Constância Flávia Junqueira Ayres, Regina Célia Bressan Queiroz de Figueiredo, Antonio Pereira-Neves

## Abstract

The nasal epithelium is an initial site for SARS-CoV-2 infection, responsible for the ongoing COVID-19 pandemic. However, the pathogenicity and morphological impact of SARS-CoV-2 on the nasopharynx cells from symptomatic patients with different viral loads remain poorly understood. Here, we investigated the ultrastructure of nasal cells obtained from individuals at distinct disease days and with high and low SARS-CoV-2 loads. Squamous and ciliated cells were the main cells observed in SARS-CoV-2 negative samples. We identified virus-like particles (VLPs) and replication organelles (RO)-like structures in the squamous cells from high viral load samples after 3- and 4-days of symptoms. Ultrastructural changes were found in those cells, such as the loss of microvilli and primary cilium, the increase of multivesicular bodies and autophagosomes, and signs of cell death. No ciliated cells were found in those samples. Squamous cells from low viral load sample after 5 days of symptoms showed few microvilli and no primary cilium. VLPs and RO-like structures were found in the ciliated cells only. No ultrastructural alterations were seen in the cells from low viral load individuals after 10- and 14-days of symptoms. Our results shed light on the ultrastructural effects of SARS-CoV-2 infection on the human nasopharyngeal cells.

## Introduction

The Coronavirus Disease 2019 (COVID-19) pandemic, due to the severe acute respiratory syndrome coronavirus 2 (SARS-CoV-2), has exerted a devastating and catastrophic social and economic crisis on global healthcare systems^1–3^. The outbreak first emerged at city of Wuhan, China, at the end of 2019 and since then it has spread fast all over the world. As of 2 July 2021, more than 184 million cases of COVID-19, with more than 3.99 million deaths, have been reported globally^4^, making it one of the deadliest pandemics in recent history. The main routes of transmission of SARS-CoV-2 involve respiratory droplets and direct contact. The clinical spectrum of COVID-19 is protean, including asymptomatic cases and a wide range of symptoms from mild flu-like to severe respiratory failure, leading to considerable morbidity and mortality^5^.

The research community and companies worldwide have working to develop vaccines and treatments to slow the pandemic and mitigate the COVID-19’s damage. Currently, some available vaccines have already been used as the main effective measure against the disease. However, this is a fragile scenario with the emergence of new SARS-CoV-2 variants, unequal access to vaccines, and transmission and deaths levels still high threatening to undo progress to date. Until to reach a high vaccination coverage, required to achieve a global herd immunity^6^, the monitoring, surveillance and control measures of disease are still needed. Thus, understanding the virus-host cell interactions is essential to develop alternative treatments, vaccines, and better diagnosis methods.

In this sense, electron microscopy studies of SARS-CoV-2 have been crucial to unravel insights into the attachment, replication, migration, transmission and pathogenicity of the virus in the hosts^7–10^. The β-coronavirus SARS-CoV-2 is an enveloped, spherical shaped, positive-sense single-stranded RNA virus, ranging about 60 to 140 nm of diameter^11^. On binding to epithelial cells of the human nasopharynx, SARS-CoV-2 starts replicating and migrating down to the airways and enters alveolar epithelial cells in the lungs^12^. SARS-CoV-2 targets cells such as nasal and bronchial epithelial cells and pneumocytes, through the viral structural spike (S) protein that binds to the angiotensin-converting enzyme 2 (ACE2) receptor^13^. The ACE2 receptors are abundant in the cardiovascular system, intestine, lungs, nasal and oral mucosa, stomach, kidneys^14–16^, and in the nasal epithelial cells^17^. The rapid replication of SARS-CoV-2 in the lungs may trigger a strong cytokines storm, leading to respiratory failure^12^.

Like other virus of Coronaviridae family, SARS-CoV-2 replicate in modified endomembrane compartments called replication organelles (RO). The RO are induced after the entry of coronaviruses into cells and comprising three membrane structures that are interconnected as well as derived to the endoplasmic reticulum (ER): convoluted membranes, double-membrane spherules, and double-membrane vesicles (DMVs). Within the RO, DMVs are the key areas where viral RNA synthesis^18,19^ takes place. Subsequently, the nucleocapsids are assembled along the ER secretory pathway and Golgi complex and the virions are released to the extracellular environment by exocytosis^20^.

Diagnosis of COVID-19 is typically made using real-time polymerase chain reaction (RT-PCR) testing via naso- or oropharyngeal swab^21^. Although RT-PCR detects the viral nucleic acid and estimates the viral load, this technique is limited to determine the SARS-CoV-2 pathogenicity during the disease course and patients with different viral loads. Morphological findings have shown that SARS-CoV-2 causes severe ultrastructural alterations, such as, cell fusion, destruction of epithelium integrity, cilium shrinking, and apoptosis^22^. However, most of those studies have been performed using in vitro cell models or cells from the lower respiratory tract^23–25^. Little is still known about the impact of SARS-CoV-2 on the epithelial cells in the nasopharynx. In addition, the scope of potential target cells, and the variance in viral tropism in the epithelium of the upper airways across patients and disease courses have not been fully determined yet. Understanding how the upper respiratory epithelium responds to SARS-CoV-2, and the role of nasal squamous cells during the viral infection and replication may be useful for future therapeutic strategies.

Here, we took the advantage of the electron microscopy to investigate and compare the ultrastructure of nasopharyngeal epithelial cells obtained from symptomatic individuals at distinct disease days and with high and low SARS-CoV-2 loads as detected by RT-qPCR. We identified the presence of virus-like particles (VLPs) and RO-like structures in the squamous epithelial cells from patients with high viral load. Ultrastructural changes were found in those cells, such as the loss of microvilli, the increase of multivesicular bodies and autophagosome-like structures, and signs of necrosis. No significant changes were seen in squamous cells from low viral load individuals, and VLPs were found in the ciliated cells only. Our data highlight insights into the morphological impact of SARS-CoV-2 infection on the human upper airway epithelium.

## Results

### Patient profile and symptomatology

The demographic and clinical profiles of patients are shown in Suppl. Table S1. The median age of patients was 45 years and 76.47% of them were woman. Few patients had underlying comorbidities, including diabetes (11.76%), chronic cardiovascular disease (17.64%) and obesity (11.76%). The most common symptoms were headache (58.82%), nasal congestion (29.41%), weakness (29.41%) and cough (29.41%).

### SARS-CoV-2 detection by RT-qPCR

Samples with Ct (cycle threshold) values ≤ 40 for viral envelope gene (E) were considered positive for SARS-CoV-2. Therefore, the viral RNA was found in 6 (35.29%) of 17 samples. Three samples exhibited Ct values < 25 for viral gene (E), indicating higher viral loads, whereas three samples showed Ct values > 30, indicating lower viral loads. In the negative samples, the Ct value for viral gene was undetermined. The RT-qPCR reaction allowed the classification of samples into three groups: (a) high viral load (n=3); (b) low viral load (n=3); and (c) negative (n=11). The high viral load group included Ct values of 18.39, 23.93 and 24.42 (SD±3.35) for viral gene (E), and 27.07, 27.76 and 29.39 (SD±1.19) for endogenous gene (RP). The low viral load group had Ct values of 38.87, 35.05 and 34.39 (SD±2.42) for viral gene (E), and 26.60, 28.97 and 27.39 (SD±1.11) for endogenous gene (RP). The Ct values of the negative group ranged from 26.25 to 30.33 (SD±1.45) for the endogenous gene (RP). Although we were not able to use a standard curve for accurate quantity detection of virus copies, the cycle threshold obtained here was used to indirectly estimate the viral load because the endogenous control did not vary greatly, as observed in our reactions with short standard deviations.

### Ultrastructural findings

The effects of SARS-CoV-2 infection on nasopharyngeal epithelial cells were observed using scanning (SEM) and transmission electron microscopy (TEM). Keratinized squamous and rounded cells, and ciliated cells were the main cells found in the samples from nasal swabs. The following groups were analyzed: (a) cells from negative individuals for SARS-CoV-2; (b) samples from patients with high SARS-CoV-2 viral load (Ct < 25) after 3 and 4 days of symptoms starting; (c) cells from individuals with low SARS-CoV-2 viral load (Ct > 30) after 5, 10 and 14 days of symptoms. The relationship among Ct values, symptoms timeline and cellular ultrastructure is shown in Suppl. Table S2.

#### Nasopharyngeal cells from negative individuals

Using SEM rounded and squamous epithelial cells exhibiting numerous microvilli were the main cell types found in the samples of patients negative for SARS-CoV-2 (Figs. 1A-B). They commonly exhibited primary cilium with the ‘9+0’ microtubular axoneme (Figs. 1B-D). Bacteria and shed vesicles were often found on their surfaces (Figs. 1B, D). Some squamous cells showed the typical furrow-type pits with few or no microvilli (Figs. 1E-F). TEM confirmed and extended those SEM results. Ciliated cells with typical motile ‘9+2’ axoneme were observed (Fig. 1G). As expected, the cytosol of squamous and rounded epithelial cells was almost all densely filled by keratin filaments (Figs. 1G-L). Shedding vesicles were frequently noticed in the extracellular environment next to the cell surface (Fig. 1H). In addition, a few organelles such as lipid droplets and multivesicular bodies (MVBs) was found in some cells (Figs. 1J-L). No VLPs particles were found.

**Figure 1.**
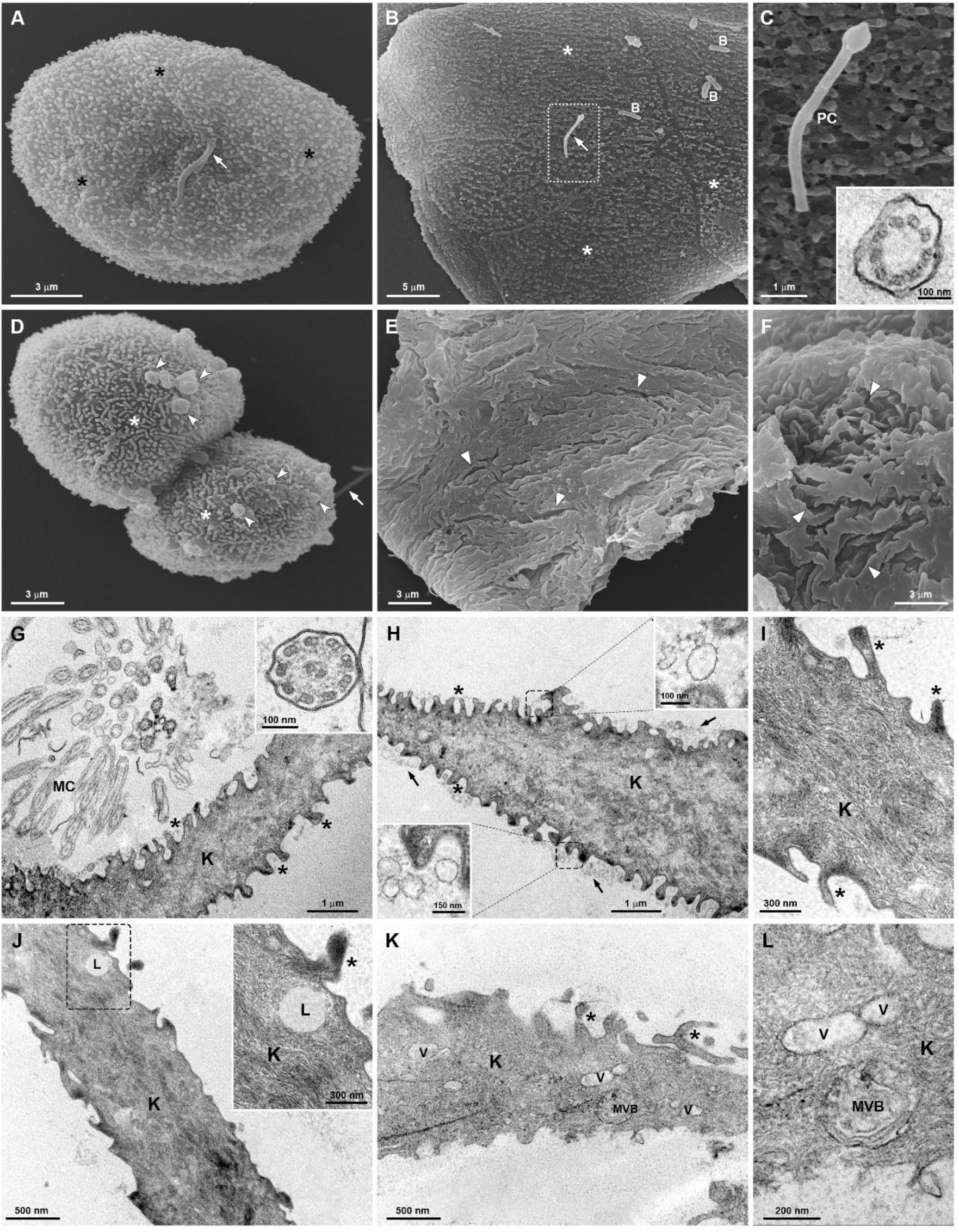
Ultrastructure of nasopharyngeal cells from SARS-CoV2 negative individuals. (**A-F**) SEM. (**A-B**) Rounded and squamous epithelial cells, respectively, with numerous microvilli (*) and primary cilium (arrow). Bacteria (B) are seen over the cell surface. (**C**) Detailed view of a primary cilium (PC). Note the ‘9+0’ microtubular axoneme by TEM (inset). (**D**) Microvesicles (arrowheads) protruding from the plasma membrane of a rounded epithelial cell. Notice the microvilli (*) and primary cilium (arrow). (**E-F**) General and detailed views, respectively, of a squamous epithelial cell with furrow-type pits (arrowheads) and no microvilli. (**G-L**) TEM. Many microvilli are seen (*) and the cytosol is densely filled by keratin filaments (K). (**G**) Motile cilia (MC) with typical ‘9+2’ axoneme (inset). (**H**) The cell exhibits some extracellular vesicles (arrows and insets) next to its surface. (**I**) Detailed view showing microvilli (*) and the cytosol with keratin filaments (K). (**J**) The cell displays a lipid droplet (L and inset). (**K-L**) General and detailed views, respectively, of a cell exhibiting some vesicles (V) and a mutivesicular body (MVB). Bars, A, D-F, 3 μm; B, 5 μm; C, G-H, 1 μm; I, 300 nm; J-K, 500 nm; L, 200 nm.

#### Nasopharyngeal cells from individuals with high SARS-CoV-2 viral load after 3 and 4 days of symptoms

Similar results were found in the samples of 3 and 4 days of symptoms. In contrast to the SARS-CoV-2 negative samples, we did not find epithelial cells displaying microvilli and primary cilium (Figs. 2-3). The squamous cells showed a smooth surface or membrane invaginations, such as furrow- and cylinder-type pits (Figs. 2A-C; Suppl. Figs. S1 A-C). Rounded cells were not found. We found numerous VLPs hetero- or homogeneously distributed over the surface of some cells (Figs. 2A-E; Figs. 3A-C; Suppl. Fig. S1). VLPs clusters were also seen covering the cell surface and in the extracellular environment (Figs. 2F-H; Figs. 3A, C; Suppl. Figs. S1 A-E). Many cells displayed cortical dense fibrils forming projections with nearby VLPs (Figs. 3D-F; Suppl. Figs. S1 C-E). A higher number of extracellular vesicles and mucus granules were observed when compared to the negative samples (Fig. 2F; Figs. 3E, G-H; Suppl. Figs. S1 F-H). We also observed some phases of the internalization of VLPs through coated pits (Figs. 3J-L). A close electron dense connection, indicative of receptor-mediated endocytosis, was observed between the VLPs and plasma membrane (Figs. 3J-K). When compared to the SARS-CoV-2 negative samples, the squamous cells exhibited profound ultrastructural alterations, such as, (a) the presence of RO-like structures containing VLPs, and (b) an increase in the amount of vesicles, MVBs, lipid droplets and autophagosome-like structures (Fig. 4). In addition, we observed in some cells several morphological alterations indicative of cell death, including (a) wrinkled cells, (b) plasma membrane disruption, (c) loss of cytoplasm content, and (d) lysed cells (Suppl. Figs. S2 A-F). Neutrophils and neutrophil extracellular traps (NETs)-like structures were also found (Suppl. Figs. S2 G-H). We did not find ciliated cells.

**Figure 2.**
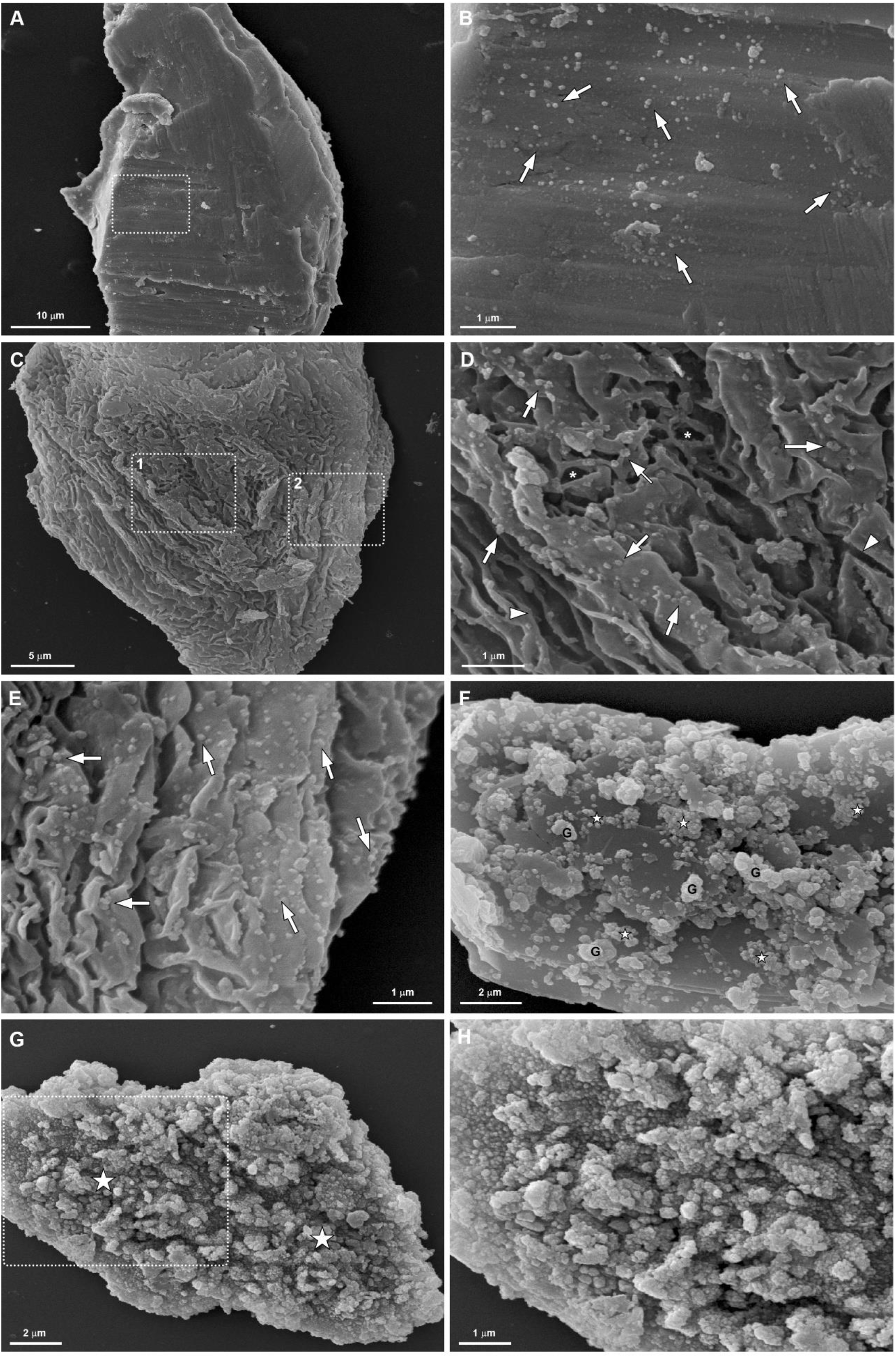
SEM of nasopharyngeal cells from individuals with high SARS-CoV-2 viral load after 3 and 4 days of symptoms. (**A-B**) General and detailed views of an epithelial cell displaying a smooth surface and some virus-like particles (arrows) heterogeneously distributed over the surface. (**C**) General view of an epithelial cell with membrane invaginations. (**D-E**) Detailed views of the regions 1 and 2, respectively, from the cell in the figure C. Observe the presence of furrow (arrowheads)- and cylinder (*)-type pits, and many virus-like particles (arrows) homogeneously distributed on the cell surface. (**F**) Clusters of virus-like particles (⋆) and some mucus granules (G) on the cell surface. (**G-H**) General and detailed views, respectively, of clusters of virus-like particles (⋆) covering all the surface of a cell. Bars, A, 10 μm; B, D-E, H, 1 μm; C, 5 μm; F-G, 2 μm.

**Figure 3.**
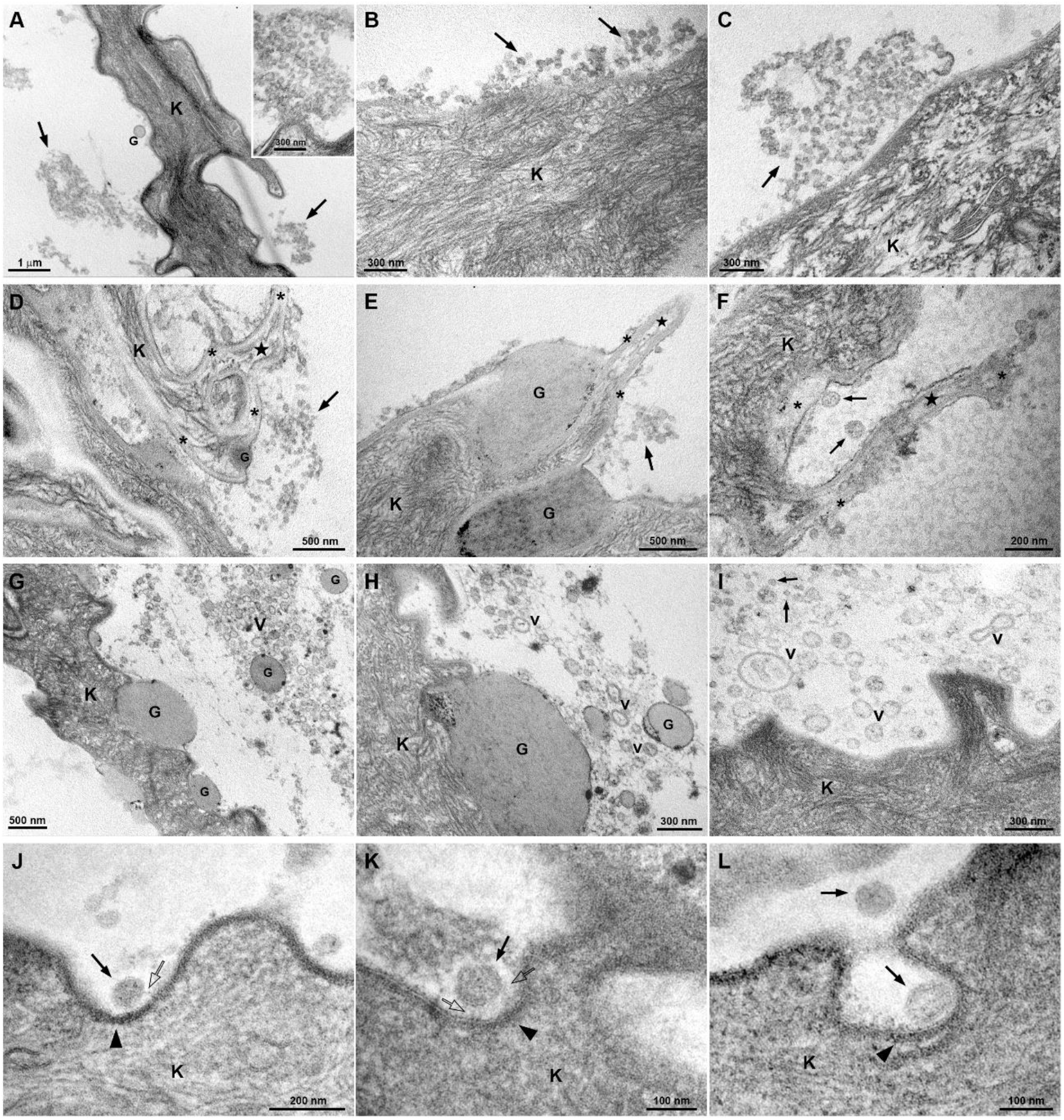
TEM of nasopharyngeal cells from individuals with high SARS-CoV-2 viral load after 3 and 4 days of symptoms. (**A**) General view of an epithelial cell. No microvilli are seen. Clusters of virus-like particles (arrows) are observed in the extracellular environment and close to the cell surface (inset). (**B-C**) Detailed views of virus-like particles (arrows) homogeneously distributed and clustered on the cell surface, respectively. (**D-F**) Surface projections (⋆) formed by cortical dense fibrils (*) with nearby virus-like particles (arrows). (**G-I**) Many mucus granules (G) and extracellular vesicles (V) are seen in the extracellular environment and close to the cell surface. (**J-L**) Phases of the internalization of virus-like particles (black arrows) through coated pits (arrowheads). Thin filamentous are seen connecting VLPs to plasma membrane (transparent arrows). K, keratin filaments. Bars, A, 1 μm; B-C, H-I, 300 nm; D-E, G; 500 nm; F, J, 200 nm; K-L, 100 nm.

**Figure 4.**
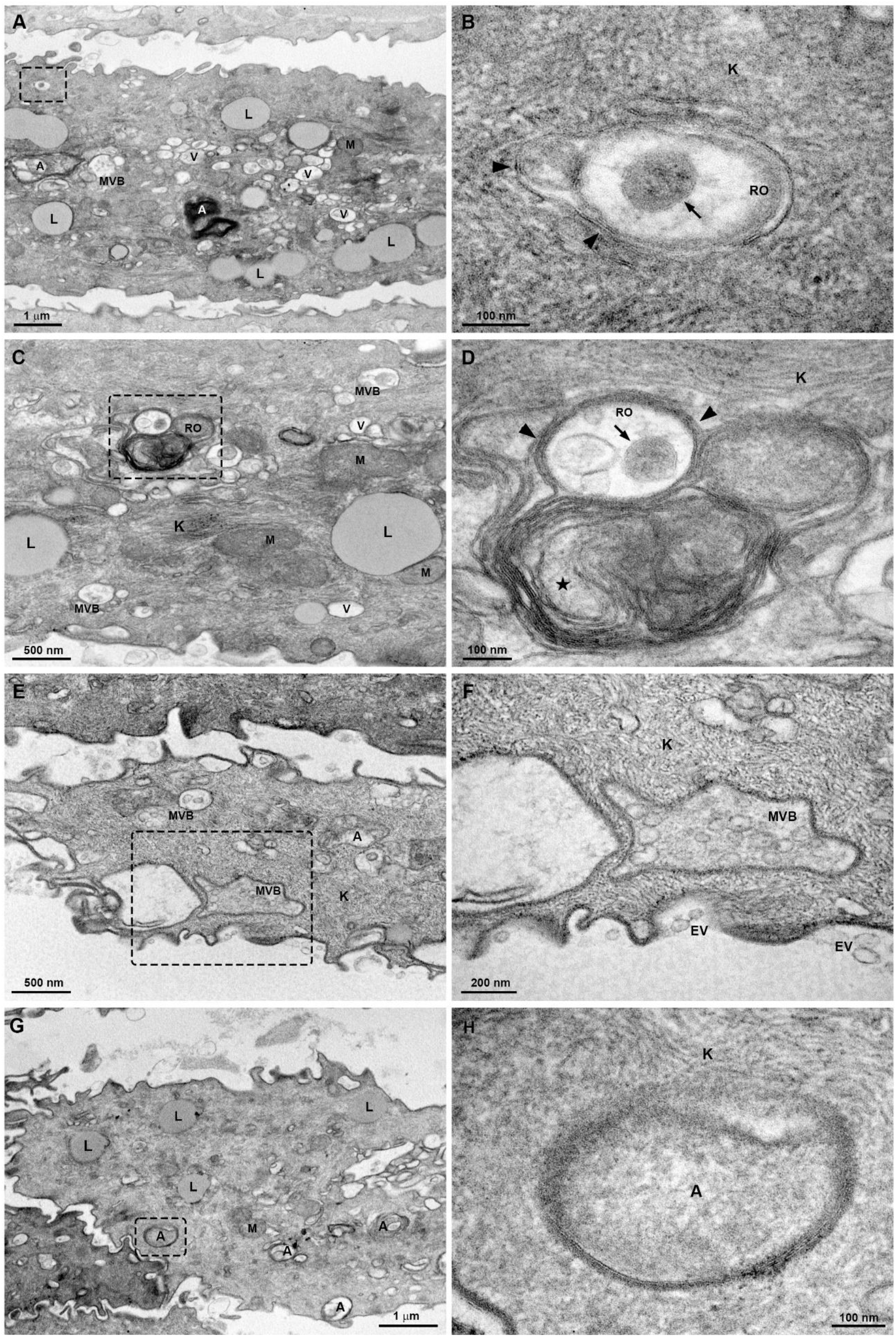
Ultrastructural alterations on nasopharyngeal cells from individuals with high SARS-CoV-2 viral load after 3 and 4 days of symptoms. General (**A, C**) and detailed (**B, D**) views of cells with replication organelles-like structures (RO). Virus-like particles (arrows) are seen surrounded by a double membrane (arrowheads) structure. Convoluted membranes (*) are also observed. Notice the presence of many vesicles (V) and lipid droplets (L) in the cytosol. (**E-F**) General and detailed views, respectively, of a cell exhibiting many multivesicular bodies (MVB). (**G-H**) General and detailed views, respectively, of a cell exhibiting many autophagosomes (A). K, keratin filaments; M, mitochondria. Bars, A, G, 1 μm; B, D, H, 100 nm; C, E, 500 nm; F, 200 nm.

#### Nasopharyngeal cells from individuals with low SARS-CoV-2 viral load

Most squamous cells from sample of 5 days of symptoms showed furrow- and cylinder-type pits without or with few microvilli (Figs. 5A-C). No primary cilium and rounded cells were found. However, the cells from patients, after 10 and 14 days of symptoms begin, exhibited morphological aspects similar to the SARS-CoV-2 negative samples, including the presence of many microvilli, primary cilium, and rounded cells (Figs. 5D-F). By TEM, similar results were found in the samples at 5, 10 and 14 days of symptoms started. Vesicles, lipid droplets, autophagosomes, MVBs and extracellular vesicles were still observed, but in an amount similar to those found in the SARS-CoV-2 negative patients (Figs. 5G-L). No VLPs or RO-like structures were found in the squamous cells. No morphological signs of cell death were seen.

**Figure 5.**
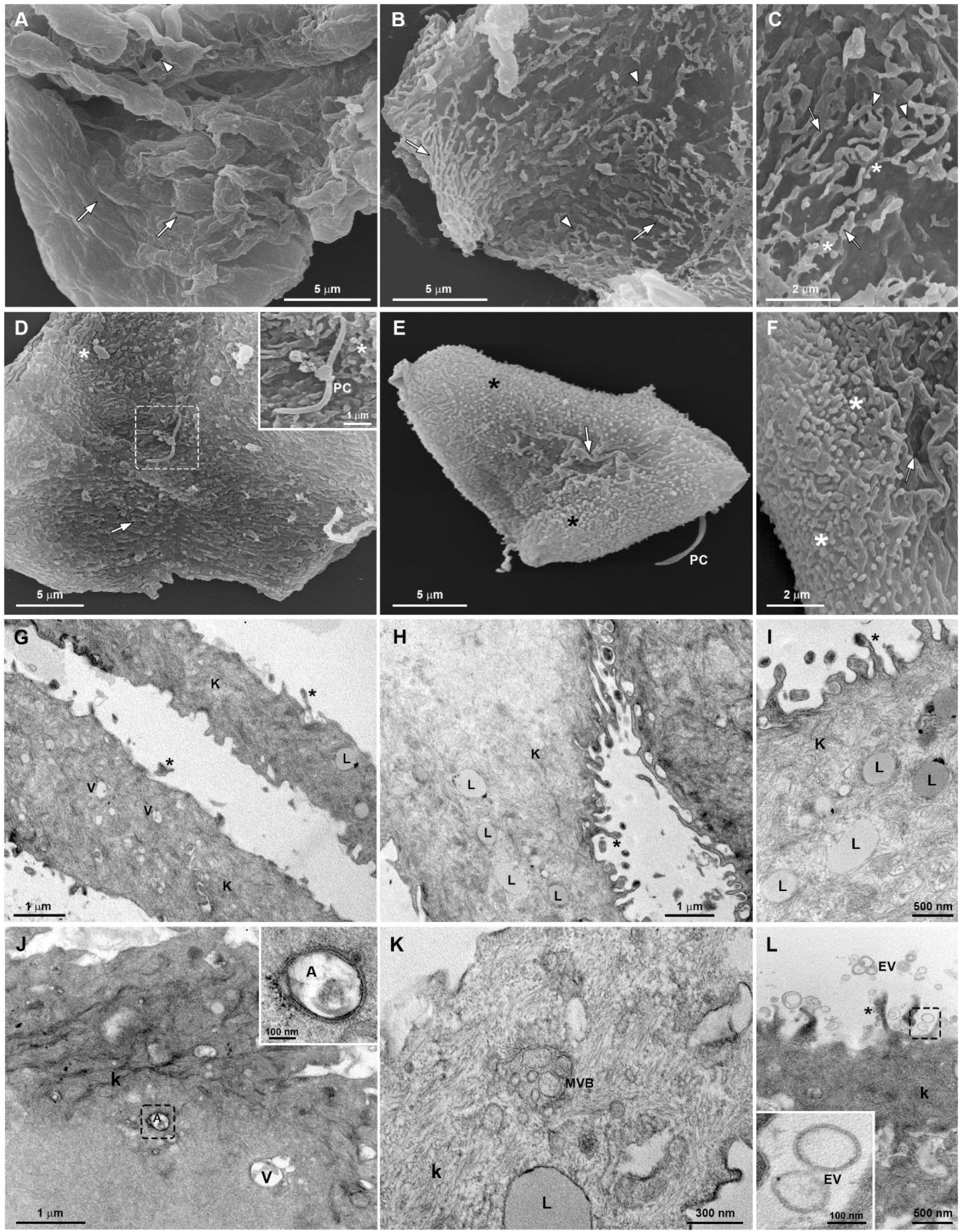
Fine structure of nasopharyngeal cells from individuals with low SARS-CoV-2 viral load. (**A-F**) SEM of samples after 5 (**A-C**), 10 and 14 (**D-F**) days of symptoms. (**A**) A squamous epithelial cell with few furrow-(arrows) and cylinder-(arrowheads) type pits and no microvilli. (**B-C**) General and detailed views, respectively, of a cell exhibiting membrane invaginations, such as furrow-(arrows) and cylinder-(arrowheads) type pits and few microvilli (*). (**D-F**) Squamous and rounded cells with many microvilli (*), primary cilium (PC) and furrow pits (arrows). (**G-L**) TEM of samples after 5 (**G**), 10 and 14 (**H-L**) days of symptoms. (**G-L**) Microvilli (*), some vesicles (V) and lipid droplets (L) are seen. (**J**) A cell exhibits an autophagosome (A and inset). (**K**) Detail of a multivesicular body. (**L**) Some extracellular vesicles (EV and inset) are noticed next to the cell surface. K, keratin filaments. Bars, A-B, D-E, 5 μm; C, F, 2 μm; G-H, J, 1μm; I, L, 500 nm; K, 300 nm.

In contrast to the high viral load samples, we found ciliated cells in all analyzed low viral load samples (Figs. 6A-B). Free VLPs and cell debris containing VLPs were observed close to these cells in the sample of 5 days of symptoms (Figs. 6C-F). Ciliated cells also exhibited RO-like structures (Figs. 6G-H).

**Figure 6.**
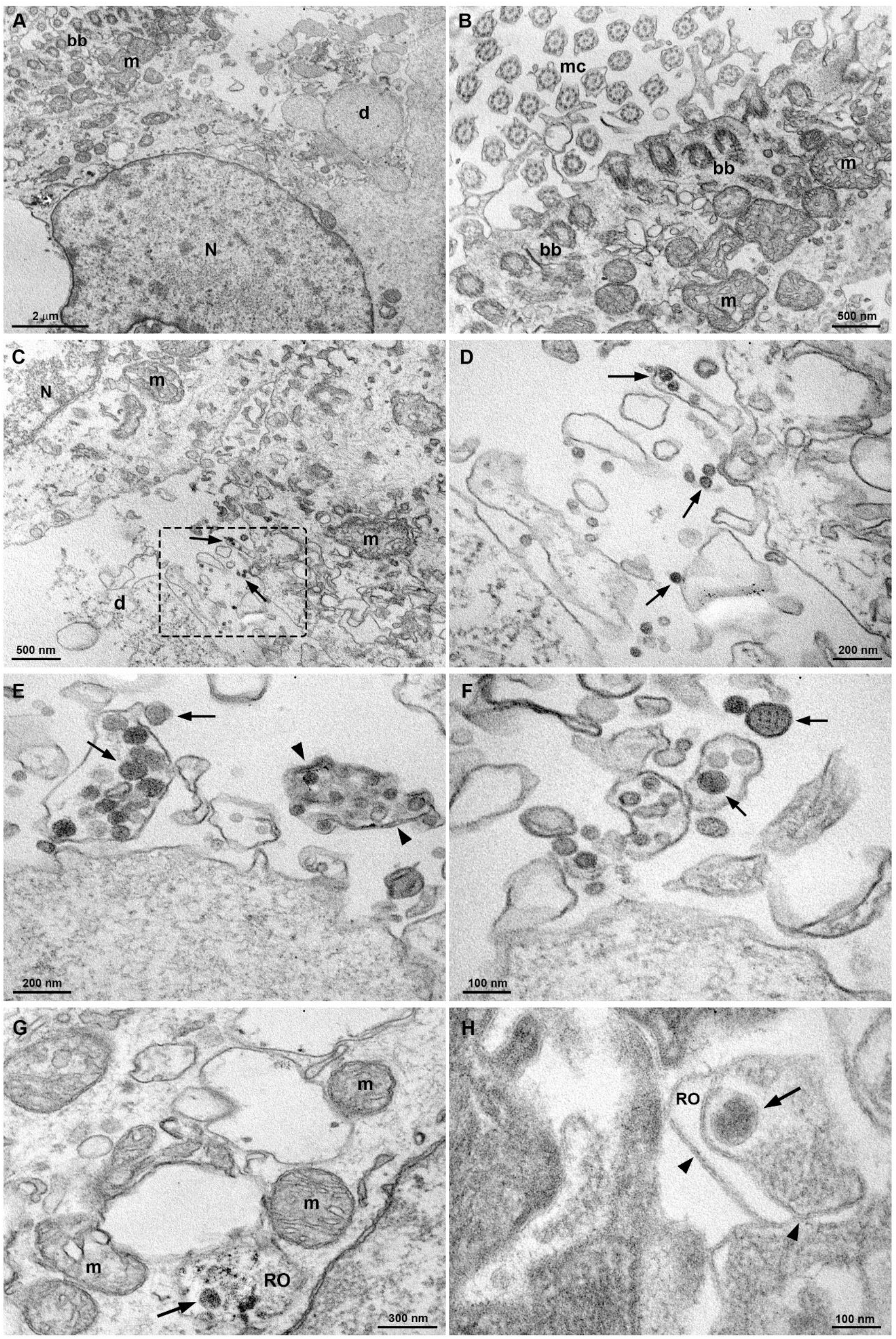
TEM of ciliated cells of samples from individuals with low SARS-CoV-2 viral load after 5 days of symptoms. (**A-B**) General and detailed views, respectively, of a ciliated cell. MC, motile cilia; BB, basal bodies; N, nucleus; D, cell debris; M, mitochondria. (**C-F**) Free virus-like particles (VLPs) and cell debris with virus-like particles (arrows) are seen close to the ciliated cells. (**E**) Note that some VLPs are surrounded by a double membrane debris (arrowheads). (**G-H**) Details of replication organelles-like structures (RO). Virus-like particles (arrows) are seen surrounded by a double membrane (arrowheads). Bars, A, 2 μm; B-C, 500 nm; D-E, 200 nm; F, H, 100 nm; G, 300 nm.

## Discussion

Although the nasopharynx cells are one of the most important gates for initial infection by SARS-CoV-2, acting as a reservoir for viral transmission and spread across the respiratory mucosa, the morphological changes provoked by COVID-19 infection on the upper airway cells from infected patients are poorly investigated. Here, we demonstrated by electron microscopy the ultrastructural effects of different SARS-CoV-2 viral loads on the human nasopharyngeal epithelial cells obtained from individuals at distinct disease days. The nasopharynx is characterized by a transitional zone between the ciliated columnar epithelium and the multilayered squamous one^26,27^. Epithelial cells located within this area are considered as an appropriate clinical sample for early virus detection by RT-PCR^28^. Usually, samples with Ct value < 40 are considered positive to respiratory viruses^29^. As expected, ciliated, rounded and squamous epithelial cells were the main cell types found by us in the samples collected by nasopharyngeal swabs.

The relationship between SARS-CoV-2 viral load and COVID-19 severity is still controversial. While some studies have associated higher viral loads with severe clinical outcomes^30–32^, others have demonstrated no correlation between Ct values and disease severity^33,34^. The SARS-CoV-2 has multicellular tropism^22^ and uses ACE2 and TMPRSS2 for viral entry^35^. In patients with COVID-19, the airway epithelial cells show approximately three-fold increase in ACE2 receptor expression, which is induced by interferon signaling by immune cells^36^. Moreover, ACE2 is highly expressed in keratinocytes^37,38^ and TMPRSS2 is abundant in squamous cells, suggesting that those cells are likely targets for SARS-CoV-2^39^. In this work, we observed a morphological pattern of nasal epithelial cells between symptomatic individuals with high and low viral loads in different days of symptoms. Our results demonstrated several ultrastructural alterations in the squamous cells from patients with high viral loads. However, few or no significant changes were found in the low viral load samples.

Among the noticeable changes, the loss of microvilli was observed in all high viral load samples and low viral load after 5 days of symptoms sample. Deinhardt-Emmer et al. reported that SARS-CoV-2 causes the loss of microvilli in monoculture cell systems and this feature was associated to dying cells^40^. Microvilli may augment the host mucosal barrier by providing an increased surface electrostatic repulsion^41^, thereby preventing invasion of inhaled pathogens and subsequent infection^42^. In this sense, the lack of apical microvilli would impair the defense barrier and could facilitate SARS-CoV-2 invasion in the upper airway cells.

Also, an interesting alteration was the absence of primary cilium in the epithelial cells from all high viral load samples and low viral load after 5 days of symptoms sample. Primary cilium is a non-motile microtubule-based organelle that serves as a signaling platform to drive cellular responses. The description and studies of primary cilium in the airway cells are rare. Jain and colleagues reported that primary cilia play a key role in airway epithelial cell differentiation and repair, suggesting that motile ciliated cells originate from primary ciliated cells^43^. Here, this structure was often found in the epithelial cells from SARS-CoV-2 negative samples and low viral loads samples after 10 and 14 days of symptoms. Primary cilium provides target sites for coronaviruses and might acts as a potential two-way street for both viral entry and exit from the cell^44^. However, the effects of viral binding on cilium function are not fully determined yet. Although studies have shown that the loss of cilia represented one of the most striking ultrastructural features in coronavirus‐infected cells^44,45^, some evidence suggest that the deciliation is delayed and temporary, occurring at three days post-infection and returning to normal several days after recovery^44,46^. Based on this, our results suggest that the absence of primary cilium could be correlated to the days of symptoms rather than the SARS-CoV-2 load.

Moreover, our data showed that squamous cells from all high viral load samples exhibited an increase in membrane invaginations, such as cylinder- and furrow-type pits and surface projections with VLPs. Those membrane alterations resemble the mechanism previously described as “cell surfing”, which is involved in viral propagation in the respiratory epithelium^23^. Some of the squamous cells showed enlarged filamentous cortical area that are similar to actin cortex. The actin polymerization serves as an additional force for membrane bending during viral budding^47^. SARS-CoV-2 exploits receptor-mediated endocytosis through interaction between its spike with host cell receptors^48^. In agreement, here, we found VLPs connected to squamous cell plasma membrane by thin filamentous structures through coated pits, which is a remarkable ultrastructure feature of receptor-mediated endocytosis.

We also found free or clustered VLPs over the cell surface and VLPs surrounded by double-membrane vesicles (DMVs), similar to RO, in the squamous cells from the high SARS-CoV-2 load samples, but not in those cells from low viral load samples, suggesting a correlation between the appearance of RO-like structures in the squamous cells and viral load. RO can work as scaffolds to anchor viral replication and transcription complexes, being essential for completion of the viral replication cycle^49^. Based on this, our data suggests that SARS-CoV-2 could infect and replicate in squamous cells; however, this might depend on a high SARS-CoV-2 load after 3 and 4 days of symptoms at least. In agreement, Deinhardt-Emmer et al. shows that epithelial cells in culture systems can be infected with high SARS-CoV-2 loads^40^.

An increase of extracellular vesicles, MVBs, autophagosomes and lipid droplets was only observed in the squamous cells from the high SARS-CoV-2 load samples. In general, those structures are involved in the secretory pathway. Virus-infected cells secrete various lipid-bound vesicles, including exosomes and microvesicles that are released into the extracellular environment to either facilitate virus propagation or regulate the immune responses^50^. The extracellular vesicles may incorporate virulence factors including viral protein and viral genetic material^50^.

MVBs and autophagosomes have been reported to be manipulated by viruses to promote their own life cycle and immune evasion^51^. The viral infection could induce the formation and persistence of autophagosomes either by generating immature autophagosomes or by mitigating their degradation^52^. The autophagosome provides a membrane-bound, protected environment to generate viral progeny, and viruses can use autophagy-generated metabolites for replication^51^. In addition, viruses can redirect autophagosomes to MVBs and hijack the secretion machinery for exocytosis^51^. Alternatively, host cells can activate the autophagic machinery to control viral infections, regulate the inflammatory response and promote antigen presentation^51^.

Viruses can directly activate lipophagy to supply mitochondria with fatty acids and maintain the high level of ATP required for viral replication^51^. It was demonstrated that lipid droplets are increased in host cells from monoculture systems and required for the formation of infectious virus particles during SARS-CoV-2 infection^53^. In agreement, our results show lipid droplets nearby to RO-like organelles and a positive correlation between the increase of lipid droplets and SARS-CoV-2 load, suggesting the participation of lipids on SARS-CoV-2 virion assembly in squamous cells from nasal cavity.

A positive correlation between cell death signs in the squamous cells and viral load was also found here. Severe ultrapathological changes induced by SARS-CoV-2 infection have been described, including, cell fusion, apoptosis, destruction of epithelium integrity, cilium shrinking and plasma membrane disruption^40,22^. In addition, the SARS-CoV-2 infection can trigger exacerbated inflammatory responses, contributing for the severity of respiratory illness. In this sense, neutrophils are the most abundant leukocytes that infiltrating tissues after viral infection^54^. Their antimicrobial activities are mediated by the release of antimicrobial peptides, phagocytosis as well as the formation of neutrophils extracellular traps (NETs). These structures are composed by DNA containing histones and granular proteins, such as elastase and myeloperoxidase^55^. The increase in the NET formation has been correlated with tissue damage in COVID-19 patients^56^. In the present study, we found neutrophils and NETs-like structures in infected individuals with high SARS-CoV-2 loads.

Here, ciliated cells were not observed in high viral load samples, but we found them in all SARS-CoV-2 negative and low viral load samples. COVID-19 has also been associated with the loss of mature ciliated cells during early SARS-CoV-2 infection in nasal mucosa followed by epithelial cell expansion, probably as compensatory repopulation mechanism for replacing the damaged ciliated epithelium^39^. Here, we were not able to determine why the ciliated cells were absent in high SARS-CoV-2 loads samples. It is still controversial whether the ciliated cells are destroyed by the virus, or the cilia are just disassembled or lacked^44,45^. SARS-CoV-2 displays a tropism for ciliated cells^22^. In this sense, in the sample with low viral load after 5 days, we found ciliated, but not squamous, cells exhibiting RO-like structures and extracellular VLPs nearby to them, indicating the cellular tropism of SARS-CoV-2 for ciliated cells in the upper respiratory tract.

Taken together, we provided insights into nasopharyngeal cell morphology from patients with COVID-19. Our data provide clues that, in addition to ciliated cells, the squamous cells of the upper respiratory tract are also a target for virus uptake and might be an important site of viral replication within the first days of illness and under a high SARS-CoV-2 load, at least. In low viral load-infected patients, ciliated cells could maintain the viral replication in the upper airway until viral clearance or tropism to other tissues. Those findings support the notion that SARS-CoV-2 is fully adapted to the human airway and the viral multicellular tropism^22^. Although a plenty of studies on SARS-CoV-2 have been performed over short-period, few have focused on the ultrastructural features of cells from infected individuals. Moreover, many of those studies have highlighted the importance of the lower airway’s cells^22,24,25^, and neglected the role of nasal epithelial cells during SARS-CoV-2 infection. One of the limitations in our work is the lack or unavailability of infected individuals with (a) high SARS-CoV-2 loads after more than 4 days of symptoms, and (b) low viral loads with days of symptoms lower than 5. Nevertheless, the results reported here should bring new insights into the impact of SARS-CoV-2 infection on the upper airway epithelial cells and have implications for the understanding of viral transmission and pathogenesis, especially during the first week of symptoms, when the virus shedding is very high^57^.

## Methods

### Patients and samples collection

This was a retrospective cohort study. Patients with signs or symptoms of respiratory illnesses and those who have had contact with people affected by COVID-19 were admitted to Oswaldo Cruz Foundation Brazil (FIOCRUZ-PE) in July 2020. A total of seventeen patients were enrolled in this study. They signed consent and filled out a form including demographic and clinical information. After that, specimens from the respiratory mucosa were collected with cotton swabs and used for SARS-CoV-2 detection by RT-qPCR and analysis by electron microscopy. All procedures were performed in accordance with WHO guidelines^58^. This work was approved by the Aggeu Magalhaes Institute Ethical Committee – CAAE 32333120.4.0000.5190.

### RT-qPCR testing for SARS-CoV-2

RNA was extracted from nasopharyngeal swab samples using Maxwell® 16 Viral Total Nucleic Acid Purification Kit (Promega, Wisconsin-USA) according to the manufacturer’s instructions. The one-step RT-qPCR assay was performed using Molecular SARS-CoV-2 (E/RP) Kit (Bio Manguinhos/Fiocruz, Rio de Janeiro, Brazil) for detecting the SARS-CoV-2 envelope gene (E) and the Ribonuclease P housekeeping control gene (RNAse P)^59^. The QuantStudio™ 5 (Thermo Scientific, USA) system was used for RT-qPCR under the following conditions: 50 ºC for 10 minutes, 95 ºC for 2 minutes, 95 ºC for 20 seconds and 45 cycles of 58 ºC for 30 seconds. Positive and negative controls were used in all plates for diagnostic. Samples were positive for COVID-19 when Ct values were ≤ 40 for gene E and ≤ 35 for endogenous control. The Ct values were used as indirect indicators of viral loads, and thus the lower the Ct value the higher the viral load.

### Electron microscopy of nasopharyngeal epithelial cells

The samples were prepared at a Biosafety level-3 Lab. The nasal swabs were washed with phosphate buffered saline (PBS) and the medium was centrifuged at 600g for 10 minutes. The cell pellets were fixed overnight in a Karnovsky solution containing 4% paraformaldehyde, 2.5% glutaraldheyde in 0.1 M cacodylate buffer, pH 7.4, for TEM and SEM preparations, as mentioned below.

#### TEM

Fixed cells were washed in 0.1 M cacodylate buffer and post-fixed in a solution containing 1% OsO4, 2 mM calcium chloride and 0.8% potassium ferricyanide in 0.1M cacodylate buffer. The samples were dehydrated in acetone series and embedded in Embed-812. Ultrathin sections were placed on 300-mesh nickel grids, stained with 5% uranyl acetate and 1% lead citrate. The samples were visualized with a FEI Tecnai G2 Spirit BioTwin transmission electron microscope, operated at 120 kV.

#### SEM

Fixed cells were washed in PBS and post-fixed in 1% OsO4. After that, the samples were dehydrated in ethanol series and submitted to critical point dryer with liquid CO2. The samples were coated with 15 nm layer of gold and then observed with a Jeol JSM-5600 electron microscope, operated at 15 kV.

## Data availability

All data are included in this paper and supplementary information.

## Author contributions

Conceived and designed the experiments: K.L.A.S., R.C.B.Q.F. and A.P.N. Performed the experiments: K.L.A.S. and L.R.S.V. Collection and manipulation of patient’s samples: M.F.B. and R.M.L.A. Analyzed the data: K.L.A.S., L.R.S.V., R.C.B.Q.F. and A.P.N. Contributed reagents/materials/tools/infrastructure: S.P.B.F., C.F.J.F., R.C.B.Q. and A.P.N. Wrote the paper: K.L.A.S., L.R.S.V. and A.P.N. All the authors were involved in reviewing and editing the manuscript. All authors read and approved the final manuscript.

## Funding

The authors received no specific funding for this work.

## Competing interests

The authors declare no competing interests.

## Additional Information

Correspondence and requests for materials should be addressed to K.L.A.S.

## Notes

### Competing Interest Statement

The authors have declared no competing interest.

