## Supplementary Information for "Ultrastructural analysis of nasopharyngeal epithelial cells from patients with SARS-CoV-2 infection"

[1]Instituto Aggeu Magalhães (IAM), Fundação Oswaldo Cruz (FIOCRUZ), Núcleo de Plataformas Tecnológicas. Av. Professor Moraes Rego, s/n - Campus da UFPE - Cidade Universitária, 50.740-465, Recife/PE – Brasil. [2]Instituto Aggeu Magalhães, Fundação Oswaldo Cruz, Departamento de Parasitologia. [3]Instituto Aggeu Magalhães, Fundação Oswaldo Cruz, Departamento de Microbiologia. [4]Instituto Aggeu Magalhães, Fundação Oswaldo Cruz, Departamento de Imunologia. [5]Instituto Aggeu Magalhães, Fundação Oswaldo Cruz, Departamento de Entomologia.

| **Features** | **Findings** |
|---|---|
| **Age (yr)** | |
| Mean | 45 |
| Range | 18-75 |
| **Sex (n [%])** | |
| Female | 13 (76.47) |
| Male | 4 (23.52) |
| **Symptoms (n [%])** | |
| Fever | 3 (17.64) |
| Cough | 5 (29.41) |
| Sore throat | 4 (23.52) |
| Dyspnea | 2 (11.76) |
| Coryza/nasal congestion | 5 (29.41) |
| Myalgia/arthralgia | 1 (5.88) |
| Headache | 10 (58.82) |
| Loss of smell | 4 (23.52) |
| Loss of taste | 2 (11.76) |
| Tiredness/weakness | 5 (29.41) |
| **Comorbidities (n [%])** | |
| Diabetes | 2 (11.76) |
| Chronic cardiovascular disease | 3 (17.64) |
| Obesity | 2 (11.76) |

**Supplementary Table S1.** Demographic and clinical profile of patients (n = 17).

| Ct value for viral gene (E) | Symptoms timeline (days) | Ultrastructural aspects |
|---|---|---|
| Undetermined | Asymptomatic | Presence of rounded, squamous and ciliated cells; presence of microvilli and primary cilium; extracellular vesicles; few MVBs and lipid droplets; absence of VLPs. |
| High viral load (Ct < 25) | 3-4 | Presence of squamous cells and neutrophils; absence of rounded and ciliated cells; RO-like structures and VLPs; loss of surface microvilli and primary cilium; cell death signs; surface projections formed by cortical dense fibrils; (+) extracelular vesicles; (+) cytosol vesicles; (+) MVBs; (+) autophagosomes; (+) lipid droplets; (+) mucus granules. |
| Low viral load (Ct > 30) | 5-14 | Presence of squamous, rounded (10 and 14 days) and ciliated cells; absence of VLPs and RO-like structures in the squamous cells; presence of VLPs in cell debris close to the ciliated cells (5 days); absence of cell death signs; few microvilli and no primary cilium (5 days); presence of microvilli and primary cilium (10 and 14 days); extracelular vesicles; (-) cytosol vesicles; (-) MVBs; (-) autophagossomes; (-) lipid droplets. |

**Supplementary Table S2.** Relationship among Ct values, symptoms timeline and cellular ultrastructure.

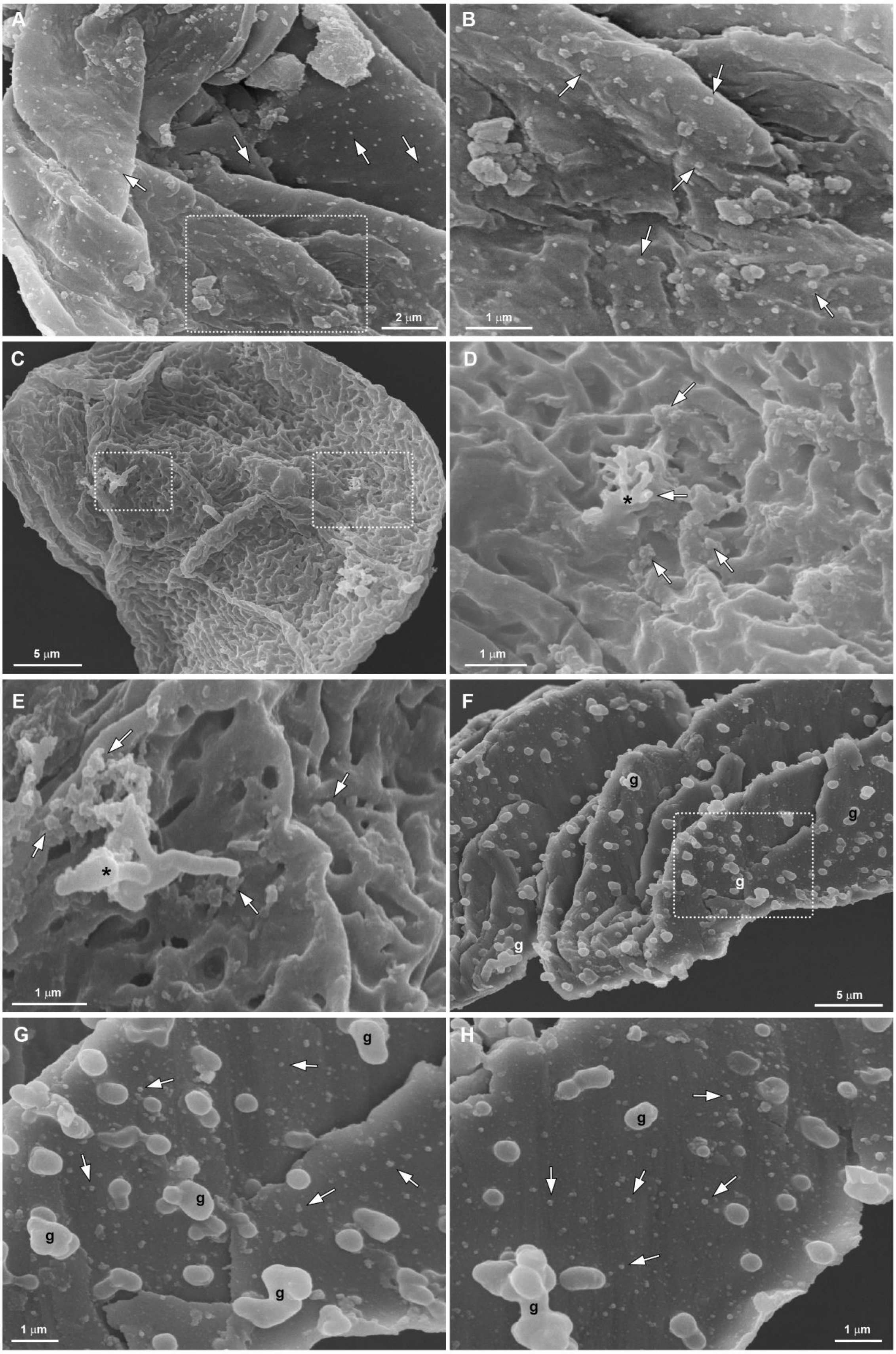


**Supplementary Figure S1. SEM of nasopharyngeal cells from individuals with high SARS-CoV-2 viral load after 3 and 4 days of symptoms.** (**A-B**) General and detailed views, respectively, of an epithelial cell displaying many virus-like particles (arrows)

homogeneously distributed on the cell surface. (**C**) General view of an epithelial cell with membrane invaginations. (**D-E**) Detailed views of the squared regions from the cell in the figure C. Notice the surface projections (*) with nearby virus-like particles and clusters (arrows). (**F-H**) General and detailed views of a cell exhibiting many mucus granules (G) and virus-like particles (arrows) on the surface. Bars, A, 2 µm; B, D-E, G-H, 1 µm; C, F, 5 µm.

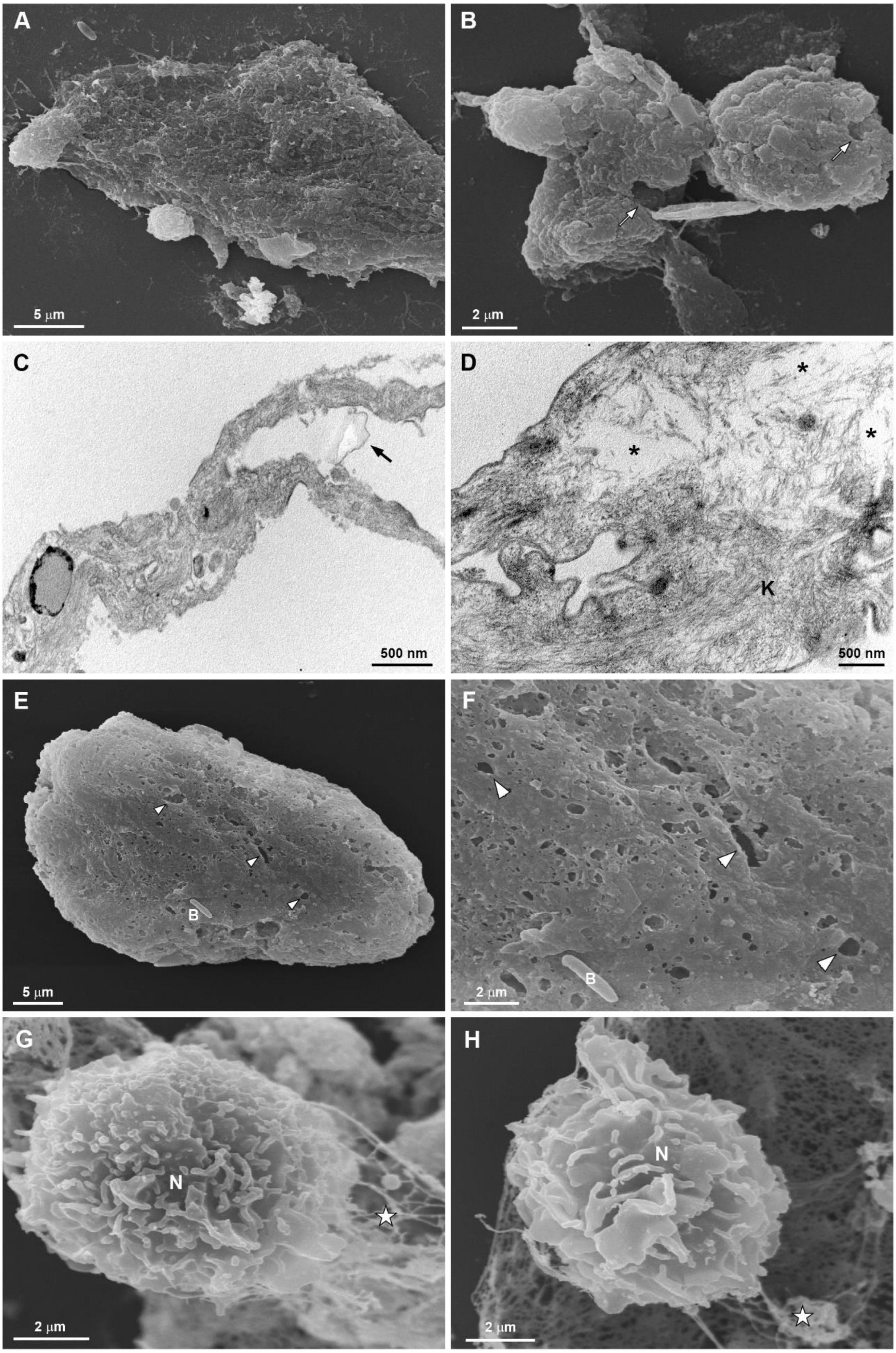


**Supplementary Figure S2. Morphological alterations indicative of cell death in the nasopharyngeal cells from individuals with high SARS-CoV-2 viral load after 3 and 4 days of symptoms.** (**A-B**) SEM of wrinkled cells with plasma membrane disruption (arrows). (**C**) TEM of a cell with plasma membrane disruption (arrow). (**D**) TEM of a cell with loss of cytoplasm content (*). K, keratin filaments. (**E-F**) General and detailed views of a lysed cell. Arrowheads indicate the surface disruption. (**G-H**) Neutrophils (N) and neutrophil extracellular traps (NETs)-like structures (★). Bars, A, E, 5 μm; B, F-H, 2 μm; C-D, 500 nm.
